# Induced pluripotent stem cell-derived human macrophages as an infection model for *Trypanosoma cruzi*

**DOI:** 10.1101/2025.03.17.643666

**Authors:** Lore Baert, Monica Cal, Thierry Doll, Matthias Müller, Pascal Mäser, Marcel Kaiser

## Abstract

Chagas disease, caused by the parasite *Trypanosoma cruzi*, affects millions of people globally. Unfortunately, the available treatment options, especially for the chronic stage of the disease, are suboptimal. Given the chronic nature of the disease and the elusive nature of the parasite, there is a high need for new and safer drugs that deliver sterile cure. Posaconazole was a promising lead in the drug discovery pipeline but ultimately failed in clinical trials due to patient relapses. This failure illustrates the need for a drug screening assay that can predict sterile cure by assessing recrudescence after treatment. Here, we used human induced pluripotent stem cell (iPSC)-derived macrophages (iMACs) as host cells for *T. cruzi*. The iMACs were highly susceptible to infection by the parasites. By combining red fluorescent protein (RFP)-expressing iMACs with mNeonGreen-expressing *T. cruzi*, we were able to monitor the dynamics of the infection through live cell imaging. The activity of the compounds benznidazole and posaconazole was consistent with the results of an established infection system using mouse primary macrophages. The post-mitotic nature of iMACs makes them suitable host cells for long-term assays needed to assess recrudescence of parasites. Moreover, their human origin, stable genetic background, and capacity for genetic modification make the iMACs excellent host cells for studying host-pathogen interaction.

**Author summary:** The parasite *Trypanosoma cruzi*, the causative agent of Chagas disease, is a global health concern affecting millions each year. Infection with *T. cruzi* can cause chronic disease, often remaining asymptomatic for decades before resulting in severe cardiac or gastro-intestinal pathologies. To date, only benznidazole and nifurtimox are used for treatment of the infection, but both drugs are suboptimal for curing the chronic stage. Posaconazole showed great promise in preclinical studies but failed to achieve sterile cure in clinical trials, causing patient relapses. These disappointing results underline the need for drug screening assays able to predict sterile cure by evaluating recrudescence post-treatment. We used human induced pluripotent stem cell derived macrophages as host cells for *T. cruzi* and testing of trypanocidal compounds. This model can be used for long-term in vitro screening assays to find new drug candidates against Chagas disease. The human origin of these cells combined with the possibility of upscaling their production make them great host cells for drug screening campaigns.

## Introduction

*Trypanosoma cruzi*, a protozoan parasite of the group of kinetoplastids, is the causative agent of Chagas disease. Up to 7 million people are currently infected, with an estimated mortality of 12,000 per year (1). While most of these cases are in Latin America, where the triatomine bugs that act as vectors are present, Chagas disease is of rising concern worldwide. This is partly because the triatomines are spreading north due to climate change, but mainly due to population migration and vector-free modes of transmission (2, 3). Chagas disease manifests in two stages: an acute phase that often stays undetected, followed by a chronic phase. The latter can stay asymptomatic for decades before causing life-threatening pathologies in cardiac and gastro-intestinal tissues (4). The only treatment options on the market, benznidazole and nifurtimox, are inefficient in treating the chronic phase of the disease and have many side effects (5). Posaconazole was a potential new drug candidate but ultimately failed to achieve sterile cure, resulting in relapses (6).

The current standard of searching for new trypanocidal compounds is based on phenotypic high content screening (HCS) of mammalian host cells infected with amastigote *T. cruzi* (7). As *T. cruzi* is capable of infecting almost any nucleated mammalian cell, various types of host cells have been used in these screens (8). These include, but are not limited to, monkey kidney epithelia Vero cells (9), peritoneal mouse macrophages (PMMs) (10), and human hepatoma Huh-7 cells (11). Our group uses PMMs in drug-screening assays to investigate sterile cure, as these cells are post-mitotic so the assay time can be elongated. Moreover, macrophages are highly infectable and sensitive to treatment (12).

However, the usage of PMMs has a range of disadvantages as well. For one, the production of the cells relies on the euthanasia of mice, which raises ethical questions. Moreover, the harvesting of PMMs is laborious, requires technical expertise, and yields rather low cell numbers. Recently, we have established a new screening assay for the related parasite *Leishmania donovani*, where PMMs are replaced by human induced pluripotent stem cell (iPSC)-derived macrophages (iMAC) (13). iMACs were readily infected with *L. donovani*, and the infection was sensitive to treatment with the reference compounds miltefosine and amphotericin B (13). This iMACs-based assay aligns with the 3R-principles and eliminates the need for lab animals for screening purposes. On top of this, it allows us to work with a more physiological human model instead. Additionally, high yields are easily obtained and the iMACs can be genetically modified, allowing us to study host-pathogen interaction also from a host-cell perspective.

Here, we show that iMACs are very useful also as host-cells for *T. cruzi* infection. Red fluorescent protein (RFP)-expressing iMACs were infected with a LucNeon-expressing *T. cruzi* STIB980 strain (10). This allowed us to follow the infection over time using live-cell imaging. Additionally, the sensitivity of the infected iMACs to the reference compounds benznidazole and posaconazole was measured and yielded results comparable to using the PMM-based assay.

## Material & Methods

### Media

*mTeSR Plus medium:* mTeSR Plus Basal Medium (pH 7.4, STEMCELL Technologies) + mTeSR Plus 5x supplement + 1% penicillin/streptomycin (Gibco). *MDM* medium: DMEM (pH 7.4, Gibco), 10% hiFBS, 1% Glutamax (Gibco), 0.055 mM β-mercaptoethanol, 1x MEM Non-Essential Amino Acids (Gibco) and 1% pen/strep. *X-Vivo 15 medium*: X-Vivo 15 (pH 7.4, Lonza), 1% Glutamax, 0.055 mM β-mercaptoethanol and 1% pen/strep. *RPMI Complete medium (iMACs):* RPMI 1640 Glutamax (pH 7.4, Gibco) supplemented with 10% hiFBS (Gibco), 1% sodium pyruvate (Gibco), 25 mM HEPES (Gibco), 0.055 mM β-mercaptoethanol (Gibco) and 40 ng/mL hm-CSF. *RPMI 1640 complete medium (PMMs & T. cruzi)*: RPMI 1640 (pH 7.4, LubioScience) + 10% hiFCS (BioConcept), 25 mM HEPES (Applichem), 24 mM sodium bicarbonate (Sigma) and 1.7 µM L-glutamine.

### Generation of RFP line

Female human WT29 iPSCs were derived from cell line AG092429 obtained from the NIA Aging Cell Repository at the Coriell Institute for Medical Research (14). To generate the red fluorescent protein (RFP)-positive WT29 iPSC line, the *RFP* gene was cloned into a PiggyBac (PB) plasmid (15) under control of the CAG promoter. For transfection, one million iPSCs, 4 µg pDNA and 1 µg transposase were mixed in 100 µL Nucleofector Solution 1 for human stem cells (Human Stem Cell Nucleofector Kit 1, #VPH-5012, Lonza). Electroporation was done using an Amaxa Nucleofector II (Lonza, program B-016). After selection with 1 μg/ml puromycin, clones were picked and analysed for homogeneous fluorescence.

### Differentiation of RFP-expressing iMACs

RFP-expressing WT29 iPSCs were grown on LN511-coated surface (BioLamina) in mTeSR Plus medium. Cells were split three times per week (1:30). The first 24 h after seeding, medium was supplemented with 10 µM Rock Inhibitor (RI) (STEMCELL Technologies). For the differentiation into iMACs, iPSCs were resuspended in mTeSR Plus medium plus 10 µM RI at 60,000 cells/mL. Using BIOFLOAT 96-well plates (FaCellitate), 100 µL of cell suspension was seeded per well. On day two, 100 µL mTeSR Plus medium containing 100 ng/mL BMP4, 40 ng/mL hSCF, and 100 ng/mL VEGF was added to the well. The day after, 100 µL of the medium was replaced with fresh mTeSR Plus medium containing 50 ng/mL BMP4, 20 ng/mL hSCF, and 50 ng/mL VEGF. On day five, the formed embryoid bodies (EBs) measured 400-700 nm in diameter. They were collected in MDM medium containing 100 ng/mL hm-CSF and 25 ng/mL IL-3. About 200 EBs were plated on a T75 flask coated with 0.1% gelatine in PBS. Following this, two-thirds of the medium were replaced with X-Vivo 15 medium containing 100 ng/mL hm-CSF and 25 ng/mL IL-3, twice per week. After three weeks, M0 iMACs started to be produced in the supernatant. The iMACs where then isolated via centrifugation (5 min, 200 g). Incubation of the cultures was done at 37 °C, 5% CO_2_.

### Isolation of PMMs

CD1 mice (Charles River Germany) were used for the harvesting of PMMs. Two days before the isolation, the peritoneal cavity was injected with 0.4 mL 2% starch solution (Sigma). After euthanising the mice, the PMMs were isolated by injecting 10 mL RPMI complete medium into the peritoneal cavity using a 25G needle, followed by extraction of the PMMs using a 22G needle. After centrifugation (10 min. 480 g), the PMMs were resuspended in RPMI 1640 complete medium, 1% antibiotic cocktail (16) and 15% RPMI medium containing growth factors obtained through seven days of cultivating LADMAC cells (mouse bone marrow cells). Next, the cell suspension was seeded in T75 flasks and incubated for three to four days in a humidified atmosphere at 37 °C, 5% CO2. The PMMs were then washed with EBSS for 30 s, and 0.05% trypsin was added for 30 s. PMMs were detached using a cell scraper and collected in RPMI complete medium.

### Cultivation of *Trypanosoma cruzi*

The *T. cruzi* STIB980 strain expressing LucNeonGreen (17) was cultivated in RPMI 1640 complete medium using mouse embryonic fibroblasts (MEF-cells) as host cells. MEF-cells were infected with an MOI of 3. After 5-7 days, trypomastigotes hatched from the cultures. The supernatant was then collected and diluted for infection of fresh MEF cells. Cultures were maintained in a humidified atmosphere at 37 °C, 5% CO_2_.

### Infection assay and imaging

iMACs or PMMs were seeded at a density of 7,500 or 10,000 cells/well in flat bottom cell culture 96-well plates (uClear, Greiner Bio-One, Cell Star) in 100 µL RPMI complete medium. Cytosine arabinoside (ara-C) was added to the iMACs at a final concentration of 2 µM. Two days after seeding, the macrophages were infected with *T. cruzi* trypomastigotes at a multiplicity of infection (MOI) of 3. After 72 h (or otherwise indicated time points) the extracellular parasites were washed away with fresh medium. After that the medium was removed from all wells once again and replaced with 100 μL medium containing a serial drug dilution of the compounds benznidazole and posaconazole. The compounds were added in a 1:3 serial drug dilution with starting concentrations of 100 µg/mL and 0.1 µg/mL, respectively. The plates were then imaged daily with the Operetta CLS (PerkinElmer). For each well, nine images were taken at 20x magnification. Infected PMMs were imaged using the EGFP channel, while infected iMACs were imaged in both the EGFP and RFP channels. Plates were always incubated at 37 °C, 5% CO_2_.

### Image analysis

Images were analysed using Harmony 4.9 software. The total number of *T. cruzi* was counted and for the infected iMACs, the total number of macrophages and the number of intracellular amastigotes was determined as well. Counting of these structures was done using automated image analysis which relied on object segmentation based on fluorescence intensity. The data were then analysed using GraphPad Prism 10.1.2 software. The data are shown as the mean ± SD. IC_50_ values were calculated by a non-linear regression curve fit, using the log(concentration) vs. the normalised response with a variable slope.

## Results

### Infection of iMACs

To develop a comparable iMAC-based assay we used our standard condition for peritoneal mouse macrophages (PMM) infection. iMACs and PMMs were infected over three days with mNeonGreen-expressing trypomastigotes at an MOI of three and monitored the infection over time (Fig. 1A). After washing away the extracellular parasites at day 3 we observed a significantly higher mNeonGreen signal in the infected iMACs (1800 ± 530 fluorescent dots) compared to the infected PMMs (280 ± 130) (Fig. 1A, B). Over the next two days, the increase in signal was similar between the two models. However, by day five, there was a noticeable increase in the signal in the iMAC infections model. We hypothesized that this could be due to cell bursting and subsequent reinfection of the cells. To better capture this dynamic, we decided to wash the infected iMACs earlier to achieve a lower infection rate. iMACs were infected for only 2, 4, and 6 hours before washing. Indeed, an infection duration of only two hours resulted in the expected low signal after three days. Utilizing RFP-expressing iMACs allowed us to calculate the infection rate and number of parasites per cell without the need for fixing and staining the cultures. By daily monitoring of the cells that had been infected for different durations, we observed that across all groups, the infection rate increased gradually in the initial days but reached high values of up to 80% after eight days of infection (Fig. 2C). Conversely, the average number of intracellular parasites increased rapidly before declining approximately five days post-infection (Fig. 2C). Imaging at various time points during the infection clearly demonstrated an increase in parasite numbers per cell until day 4, with a subsequent decrease and dissemination of the infection by day 8 (Fig. 2D). This observation supports the hypothesis that infected iMACs burst under high infection levels, leading to the reinfection of other iMACs. The bursting of iMACs was further confirmed through live imaging of RFP-expressing iMACs (Fig. 2E). Altogether, RFP-positive iMACs enable real-time monitoring of parasite infection and open new avenues for research.

**Fig. 1:**
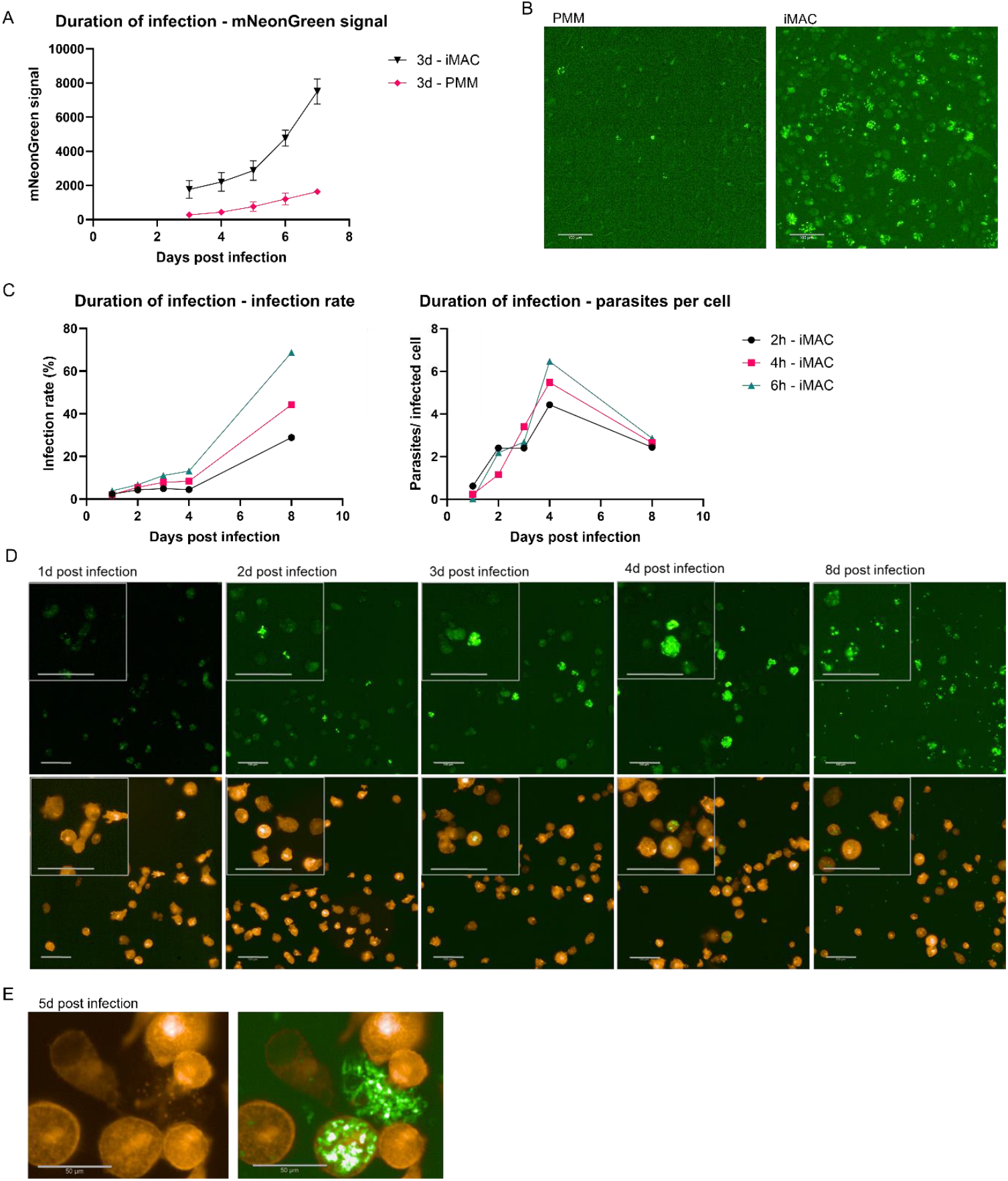
iMACs as host cells for mNeonGreen expressing *Trypanosoma cruzi*. A) iMACs and PMMs were infected with *T. cruzi* at MOI 3 and extracellular parasites were washed 3 days after infection. The infection was followed over time and measured as the number of positive mNeonGreen expressing dots. All values were corrected by subtracting the signal from negative, uninfected controls. B) PMMs and iMACs were infected at an MOI of 3. Three days after infection, the cells were washed and imaged. Representative images are shown. Pictures were taken using 20x magnification. Scale bar = 100 µm. C) iMACs were infected with *T. cruzi* at MOI 3 and extracellular parasites were washed away 2 h, 4 h or 6 h after infection. The infection was followed over time and measured as the infection rate (left) and number of parasites per infected cell (right). All values were corrected by subtracting the signal from negative, uninfected controls. D) iMACs were infected at an MOI of 3. Six hours after infection, the cells were washed and imaged. Representative images are shown. Pictures were taken using 20x magnification. Scale bar = 100 µm. E) About five days after infection, highly infected cells burst and released trypomastigotes. The bursting of the iMACs could be observed using RFP-expressing iMACs as host cells. Pictures were taken using 20x magnification. Left = RFP signal, right = overlay of RFP and GFP signal. Scale bar = 50 µm.

**Fig. 2:**
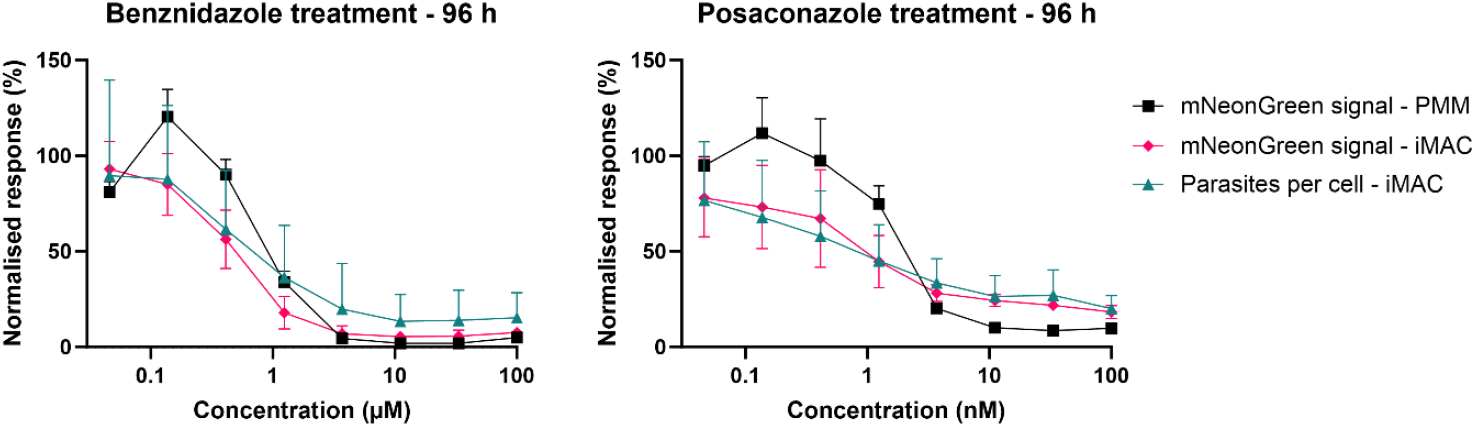
Drug-response for benznidazole and posaconazole. iMACs and PMMs infected with *T. cruzi* were treated with benznidazole and posaconazole for 96 h. The response was measured as the total mNeonGreen signal and the number of parasites per 100 cells. The data corrected by subtracting the negative control and normalised by a positive control.

### Sensitivity of iMACs to benznidazole and posaconazole

Next, we tested the sensitivity of T. cruzi in iMACs to the compounds benznidazole and posaconazole and compared the assay to the classical model using PMM. To be as close as possible to the standard assay we were using the same protocols for both cell types. Three days after infecting the iMACs with an MOI of 3, the cells were washed and the compounds were added at eight concentrations in a 3x dilution series, starting from 100 and 0.1 µg/mL, respectively. A washout of the drugs was performed after 96 h and the IC_50_ values were calculated at this time.

When looking at the overall mNeonGreen signal, we observed a comparable dose-response relationship when using iMACs or PMMs as host cells (Fig. 2). For benznidazole, we calculated IC_50_ values of 0.47 [0.40 - 0.56] and 0.98 [0.72 - 1.3] µg/mL for iMACs and PMMs, respectively, treated for 96 h. For posaconazole, the IC_50_ values for iMACs (1.0 [6.5 - 1.7] ng/mL) and PMMs (2.1 [1.5 - 2.8] ng/mL) were also comparable. (Table 1). For the treated parasites in the iMAC infection model, we also calculated the number of parasites per infected cell based on the GFP signal in the RFP labelled cell body. The IC_50_ values calculated based on the number of parasites per 100 cells were comparable to the mNeonGreen-based measurements (0.80 [0.38 - 1.7] µg/mL and 0.94 [0.39 - 2.1] ng/mL for benznidazole and posaconazole) (Fig. 2; Table 1). Based on these results, we conclude there is a comparable sensitivity of the iMAC and PMM infection model to drug treatment.

**Table 1:** IC_50_ values for treatment of iMACs and PMMs with benznidazole and posaconazole. IC_50_ values were calculated based on the total number of mNeonGreen dots (mNG) and the number of parasites per 100 cells (PpC). Calculation was done using non-linear regression in GraphPad prism. All values. 95% confidence intervals (CI) are given.

|  | PMM-mNG | iMAC-mNG | iMAC-PpC |
| --- | --- | --- | --- |
| *Benznidazole (µg/mL)* |  |  |  |
| **IC<sub>50</sub>** | 0.98 | 0.47 | 0.80 |
| **95% CI** | 0.72 - 1.3 | 0.40 - 0.56 | 0.38 – 1.7 |
| *Posaconazole (ng/mL)* |  |  |  |
| **IC<sub>50</sub>** | 2.1 | 1.0 | 0.94 |
| **95% CI** | 1.5 - 2.8 | 0.65 – 1.7 | 0.39 – 2.1 |

Next, long-term culture of infected iMACs treated with benznidazole and posaconazole was performed. Utilizing live imaging, the cells were monitored over 408 hours (17 days) (Fig. 3, Fig. S1). Up to 96 hours post-treatment, the iMACs appeared healthy and viable. However, at later time points, a decrease in viable iMACs was observed in wells that were untreated or treated with low drug concentrations, as *T. cruzi* growth led to significant cell death due to replicating parasites and bursting host cells (Fig. 3). At higher compound concentrations, the iMACs remained healthy but in lower numbers compared to the untreated negative control populations. Therefore, for time points beyond 96 hours, drug response analysis was limited to the highest benznidazole and posaconazole concentrations for the number of parasites per infected cell. These data suggest that optimisation of the assay may allow detection of regrowth of surviving parasites post-drug removal.

**Fig. 3:**
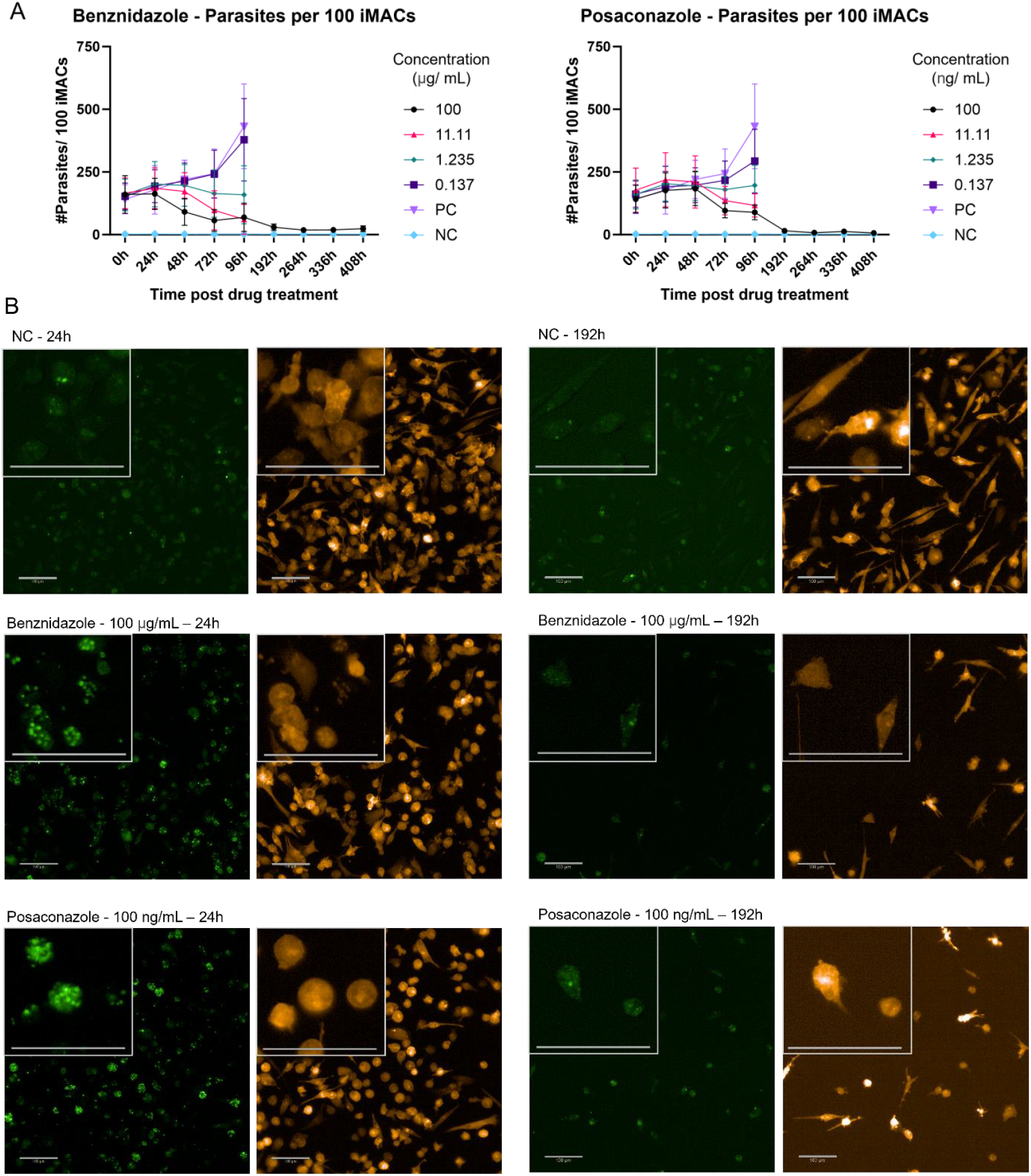
Long-term culture of infected iMACs treated with benznidazole and posaconazole. iMACs infected with *T. cruzi* were treated with benznidazole and posaconazole over 408 h (96 h drug pressure and 312 h recovery). A) The response was measured the number of parasites per 100 cells. A negative, uninfected control and a positive, infected but untreated control were included. B) Representative pictures of iMACs taken 24 h and 192 h after treatment for a negative, uninfected control (NC) and cultures treated with the highest concentrations of benznidazole and posaconazole. Pictures were taken in the EGFP (left) and RFP (right) channel at 20x magnification. Scalebar = 100 µm.

## Discussion

*Trypanosoma cruzi* parasites can infect virtually any mammalian nucleated cell, resulting in many different host cells being used in drug screening assays (8, 18). Currently, most assays used for the screening of antitrypanosomal compounds have an endpoint 72-96 h post treatment. However, this short timeframe does not suffice to assess sterile cidality of a compound, which is especially relevant considering the failure of posaconazole in phase II clinical trials (6). Therefore, our group uses a drug washout assay where the infection is followed for an additional week after the end of treatment. To allow for this, PMMs have been used as host cells as these cells are not dividing and therefore do not overgrow (12). However, the use of PMMs comes with its own drawbacks concerning ethical questions, non-human origin, limited yields, and laborious isolation protocol. Based on previous work on *Leishmania donovani* (13), we have developed a new long-term assay for *T. cruzi* using human iPSC-derived macrophages as host cells.

Using automated image analysis, we followed infection of iMACs over time during a 96 h treatment with benznidazole and posaconazole and for an additional 13 days after drug washout. The activity of the compounds was similar when using PMMs or iMACs as host cells for *T. cruzi*. However, using the same infection parameters (three-day infection with an MOI of 3) resulted in a considerably higher infection rate in iMACs compared to PMMs. The usage of RFP-expressing iMACs makes it easier to follow the infection rate and number of intracellular parasites over time, which is usually only done after fixation and staining when using non-fluorescent host cells. The RFP-expressing iMACs can therefore be used to assess pharmacodynamics *in vitro* as well as host-pathogen interaction, such as the effect on the timing of parasite release.

A disadvantage of using an automated readout based on image analysis is the inaccuracies in the analysis combined with background noise. This makes it difficult to determine whether sterile cure is achieved. Moreover, fluorescence is suboptimal to determine cidality because even a dead cell can still give a signal. To be certain of sterile cure, longer assays need to be used to assess recrudescence of the parasites after drug wash-out. Treatment periods of 16 days followed by 60 days of washout have been proposed (19). Achieving these longer time frames is difficult as we observed considerable iMAC death due to infection. This causes the loss of untreated control populations and iMACs treated at low drug concentrations after 192 h. But also in populations treated with higher drug concentrations there was a loss of cells. Optimisation of infection time and MOI could ameliorate this issue. As iMACs are highly susceptible to infection with *T. cruzi*, washing away the extracellular parasites after a few hours of infection instead of three days can provide infection rates high enough for drug screening while allowing for longer assay times. Another option is adding extra iMACs at different time point post infection, such as after wash-out of the compound. This would replenish the host cells and give viable parasites the chance to reinfect and multiply.

Macrophages are among the first host cells to be infected in Chagas disease and serve an important immunomodulatory role (20, 21). iMACs can easily be genetically modified, which is valuable for studying host-pathogen interaction (22). However, other host cell types also play a crucial role in pathology. Chronic infection of Chagas disease often results in severe cardiomyopathy. The use of human iPSC-derived cardiomyocytes has already been reported for drug screening in *T. cruzi* (23, 24). Gastro-intestinal pathology is also a common outcome of the infection, and the gastro-intestinal tract is possibly an important site for recrudescence (25). Therefore, cell lines such as iPSC-derived enterocytes could be interesting host cells for screening purposes as well (26). The host cell type used for screening can greatly influence the outcome (18). Thus, screening compounds *in vitro* against a combination of iPSC-derived host cell types can bring forth more potent drug candidates for further testing. The possibility to generate these various host cell types from a single iPSC line is a huge advantage. This allows for testing compounds against different cell types with the same genetic background, making the assays more robust and reproducible.

In conclusion, iPSC-derived macrophages are a promising new tool for screening of antitrypanosomal compounds and studying host-pathogen interaction. These cells are readily infected with *T. cruzi*, allow intracellular replication and re-infection, and their post-mitotic state allows for drug wash-out assays with much longer incubation times as compared to standard 96 h assays. The comparable efficacy of benznidazole and posaconazole in both system makes the iMACs a good alternative to PMMs. Due to their potential for upscaling, limited ethical issues, human origin and stable genetic background, we believe the iMAC are even preferable.

## Supplementary figure

**Fig. S1: Drug-response for benznidazole and posaconazole** iMACs and PMMs infected with *T. cruzi* were treated with benznidazole and posaconazole over 408 h (96 h drug pressure and 312 h recovery). The response was measured as the total mNeonGreen signal and the number of parasites per 100 cells. A negative, uninfected control and a positive, infected but untreated control were included. The mNeonGreen signal is presented as percentage, corrected by subtracting the negative control and normalised by a positive control. Starting at 192 h, only the highest drug concentrations are shown as massive cell death due to parasite infection occurred at lower concentrations and in the positive, untreated control.

